## Supplemental Info for "Correcting gradient-based interpretations of deep neural networks for genomics"

**Supplementary Table 1.** ENCODE ChIP-seq details. Ten representative TF ChIP-seq experiments in GM12878 cell line and a DNase-seq experiment (File accession: ENCFF235KUD) for the same cell line were downloaded from ENCODE. Table shows ENCODE file accession codes for all transcription factor proteins.

| PROTEIN | ENCODE FILE ACCESSION | CELL LINE |
|---|---|---|
| CTCF | ENCFF710VEH | GM12878 |
| MAX | ENCFF083KVY | GM12878 |
| ATF2 | ENCFF127GYQ | GM12878 |
| ARID3 | ENCFF027VZK | GM12878 |
| BACH1 | ENCFF012JXJ | GM12878 |
| GABPA | ENCFF116EXQ | GM12878 |
| ZNF24 | ENCFF103OOV | GM12878 |
| ELK1 | ENCFF556JBS | GM12878 |
| SRF | ENCFF909FRA | GM12878 |
| REST | ENCFF677KJB | GM12878 |

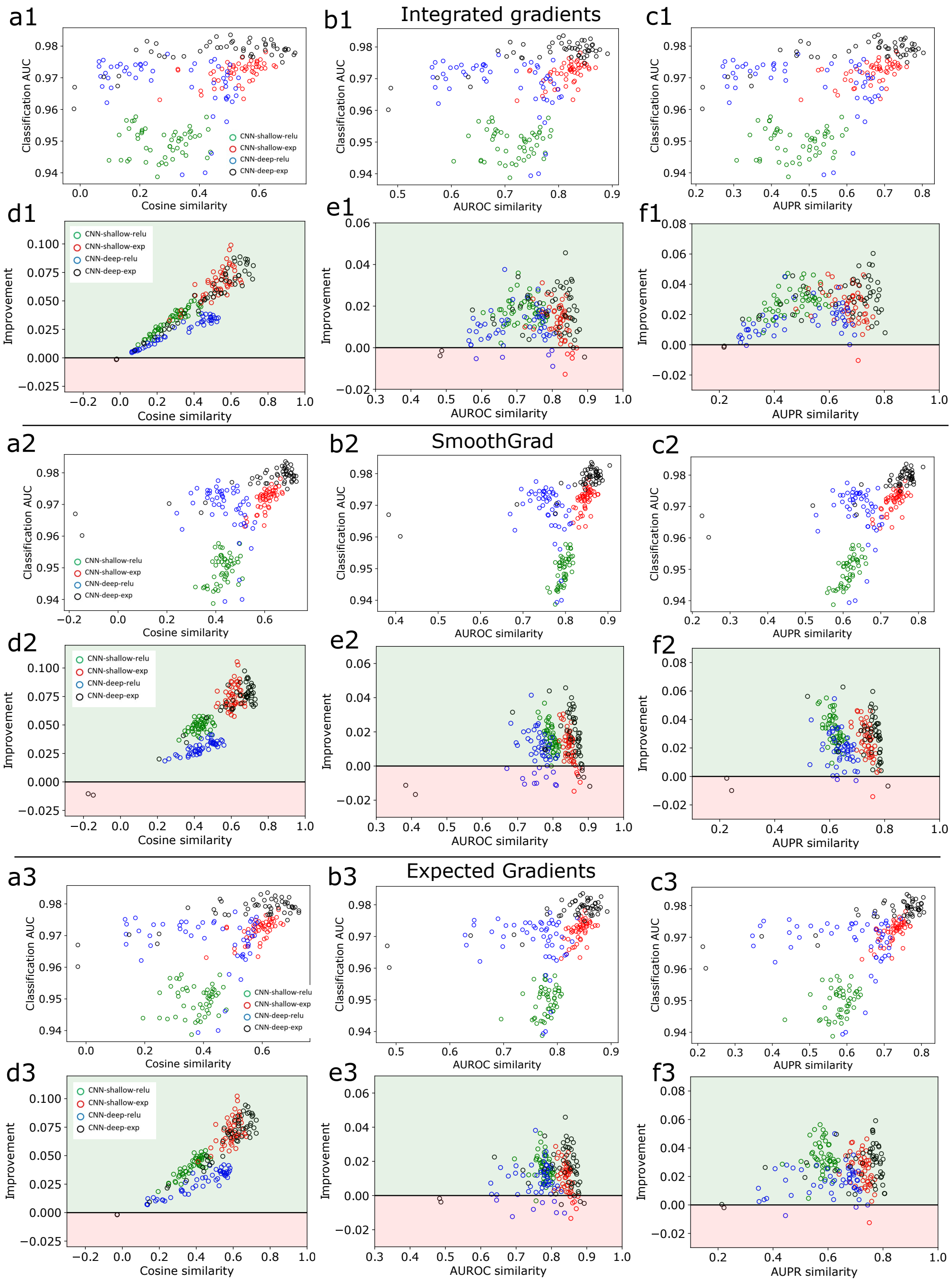


**Supplementary Figure 1.** Synthetic data performance comparison across attribution methods. Scatter plot of the classification performance according to the area-under the receiver operating characteristic curve (AUC) versus the interpretability performance measured by different similarity metrics (shown in a different column) for integrated gradients (a1-c1), SmoothGrad (a2-c2) and expected gradients (a3-c3). Interpretability improvement for integrated gradients (d1-f1), SmoothGrad (d2-f2) and expected gradients (d3-f3) for different similarity metrics when applying gradient correction. Improvement represents the difference in the similarity score after the correction minus before the correction. Green region highlights a positive improvement; light red is the region where the change in similarity score is worse. Each point represents 1 of 50 runs with a different random initialization for each model (shown in a different color).

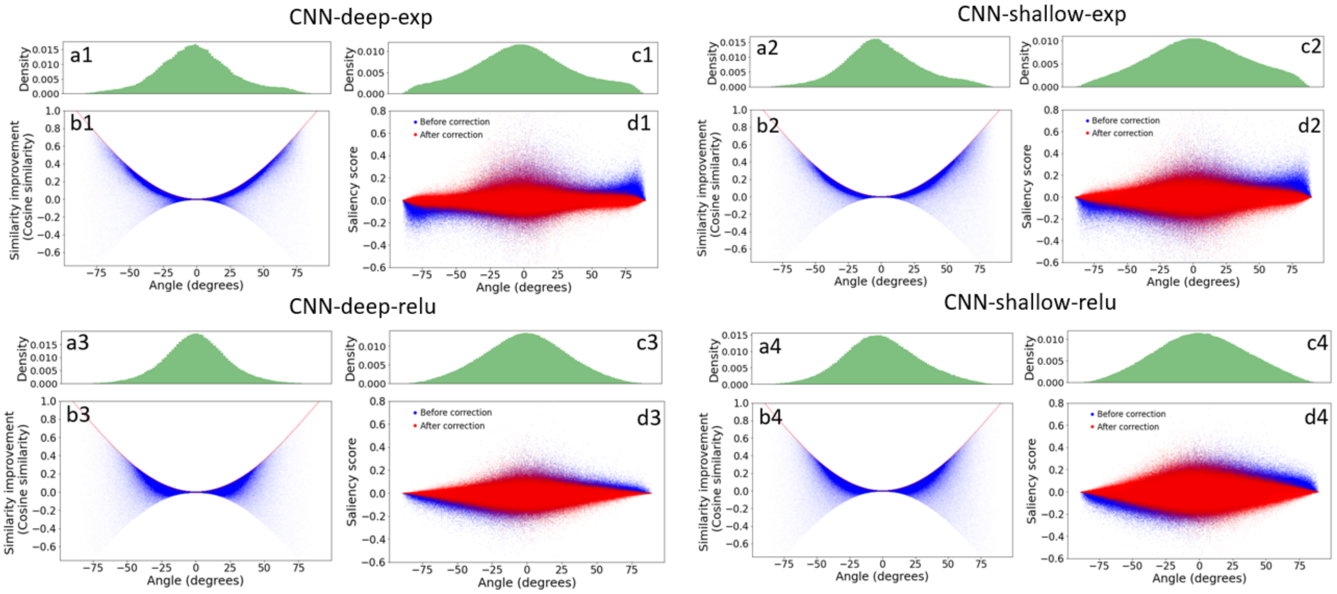


**Supplementary Figure 2.** Understanding gradient correction using synthetic data. Analysis of gradients at different angles for CNN-deep-exp (top left), CNN-shallow-exp (top right), CNN-deep-relu (bottom left) and CNN-shallow-relu (bottom right). (a, c) Probability density of input gradient angles for positions where ground truth motifs are embedded (a) and other background positions (c). (b) Scatter plot of attribution score improvements based on cosine similarity (after correction minus before correction) versus the gradient angles for ground truth positions. Red line indicates the theoretical limit for a correction, i.e. $1-\cos(\text{angle})$. (d) Scatter plot of saliency scores versus gradient angles before (blue) and after (red) correction for background positions (i.e. positions without any ground truth motifs). (b,d) Each dot represents a different position in each sequence across the test set.

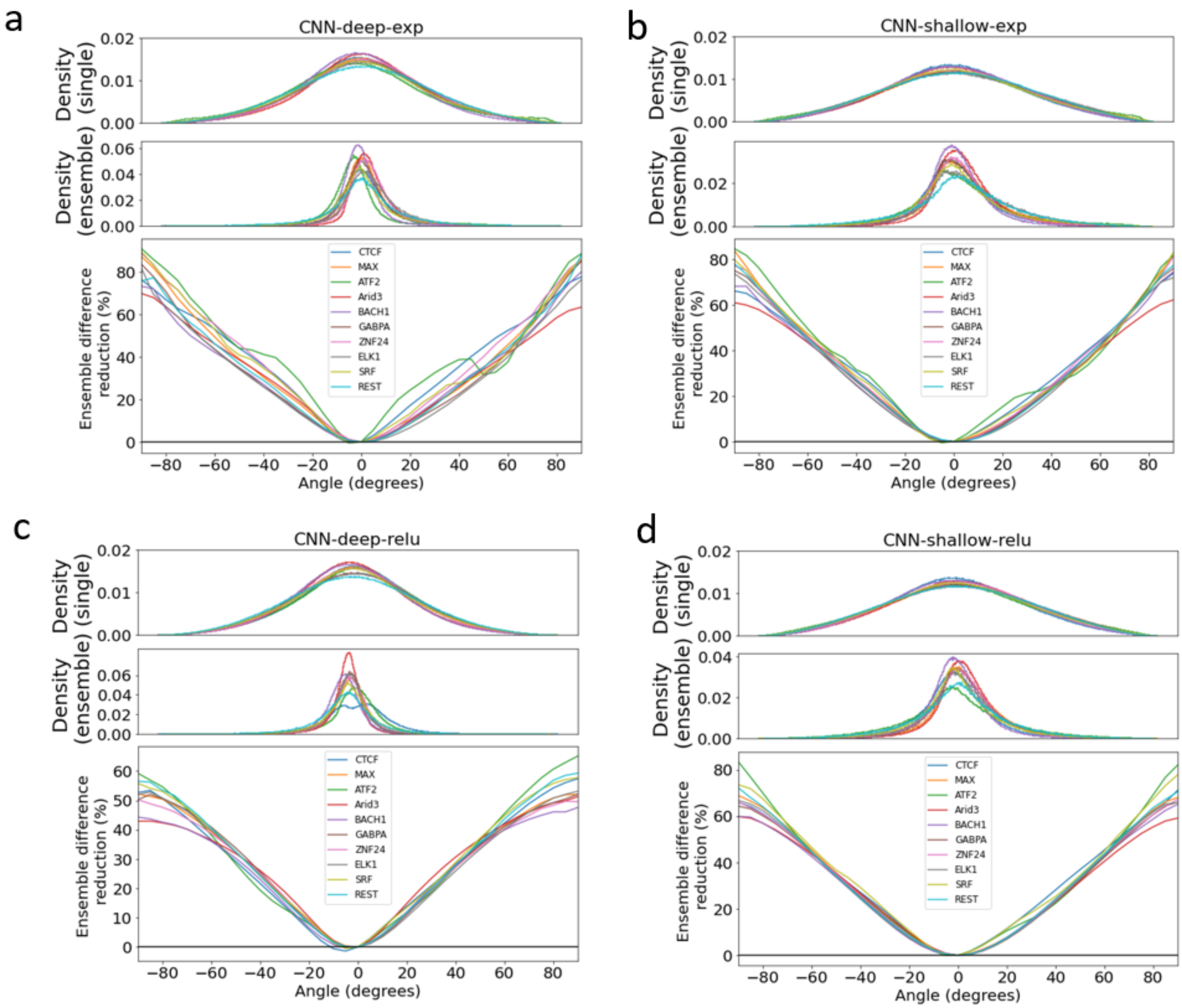


**Supplementary Figure 3.** Understanding gradient correction using ChIP-seq data. Angle-noise analysis for (a) CNN-deep-exp, (b) CNN-shallow-exp, (c) CNN-deep-relu and (d) CNN-shallow-relu for 10 different ChIP-seq proteins. (Top and middle rows in a-d) Probability density of input gradient angles. (Bottom rows in a-d) Improvement in saliency maps measured as a percentage decrease of the L2-difference between single-model saliency vector at a given position with corresponding ensemble saliency vectors serving as ground truth. Each line shows averaged results across all positions at a given angle for a different protein (shown in a different color).

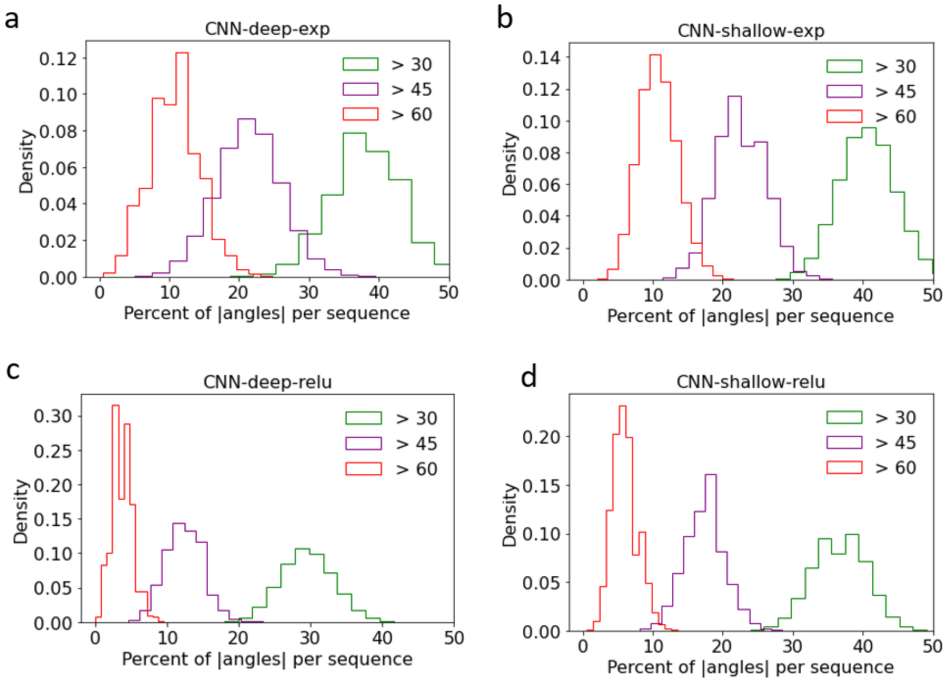


**Supplementary Figure 4.** Gradient angle analysis from CNNs trained on synthetic data. Distribution of large gradient angles for CNN-deep-exp, CNN-shallow-exp, CNN-deep-relu and CNN-shallow-relu. Histogram of the percentage of positions in a sequence with a gradient angles larger than various thresholds for each CNN model.

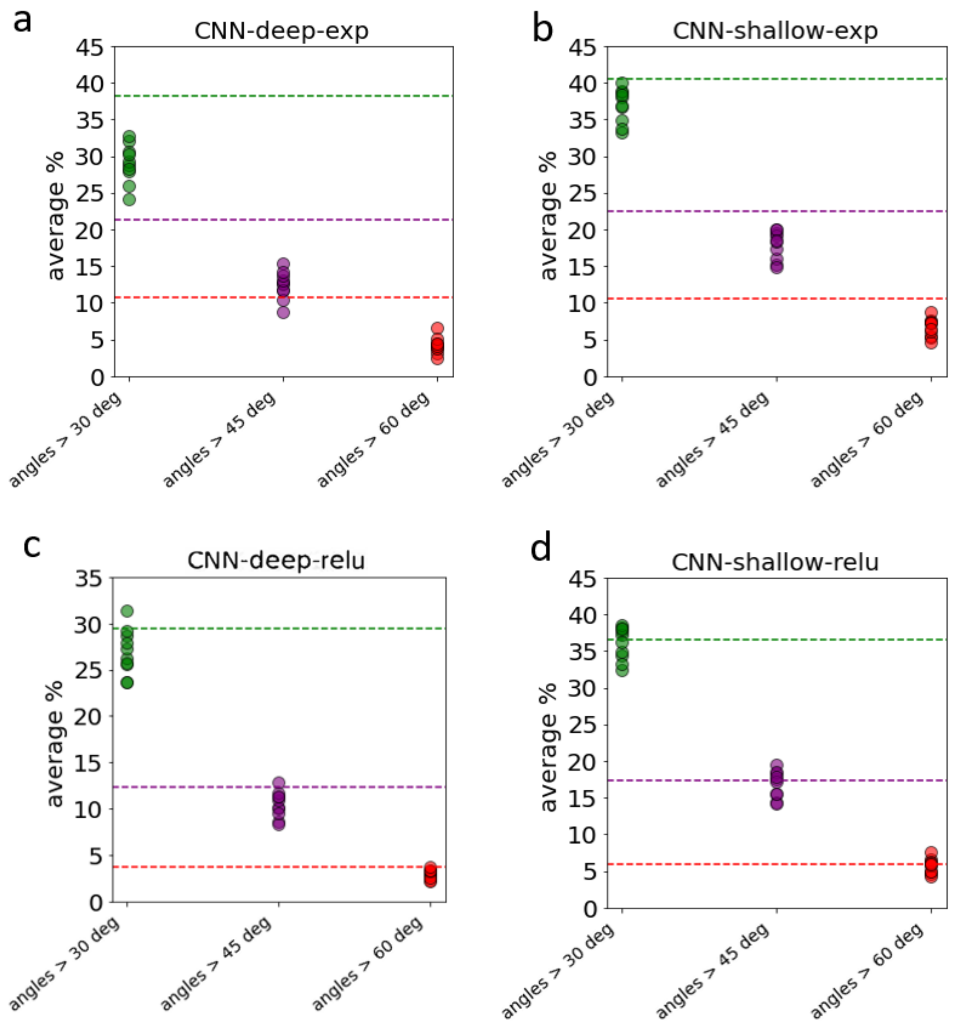


**Supplementary Figure 5.** Gradient angle analysis from CNNs trained on ChIP-seq data. Scatter plots of the average percentage of positions in a sequence with a gradient angle larger than various thresholds for CNN-deep-exp, CNN-shallow-exp, CNN-deep-relu and CNN-shallow-relu across 10 TF ChIP-seq experiments. Each point represents the results for a different ChIP-seq experiment. For comparison, horizontal dashed lines indicate the mean value from synthetic experiments using the corresponding models.

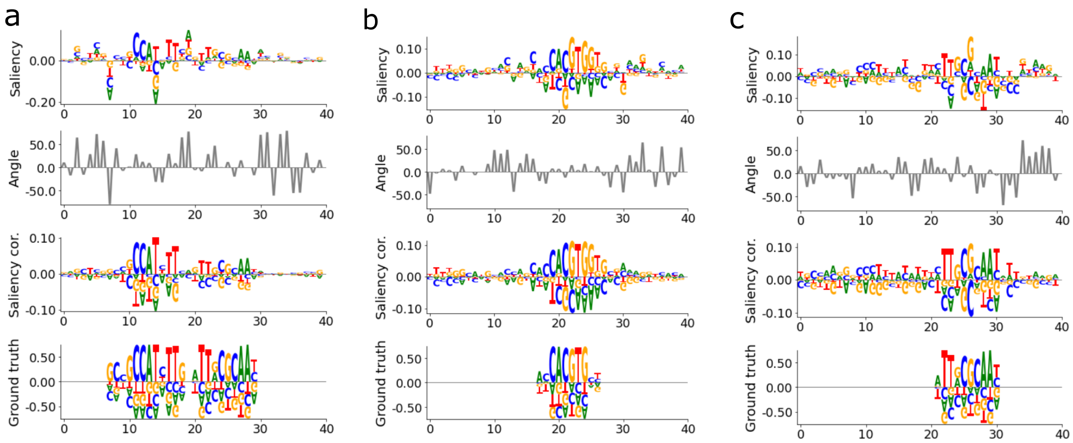


**Supplementary Figure 6.** Saliency map comparison for synthetic data. Comparison of saliency maps before and after correction for synthetic data using (a) CNN-shallow-exp, (b) CNN-deep-relu and (c) CNN-shallow-relu. (a-c) Representative patches from positive label sequences that show a sequence logo of the saliency scores at each position (top row), a plot of the angles between gradients and the simplex at each position (second row), a sequence logo of the corrected saliency scores (third row), and a sequence logo of the ground truth (bottom row). Sequence logos for ground truth are subtracted by 0.25 prior to plotting to remove uninformative positions.

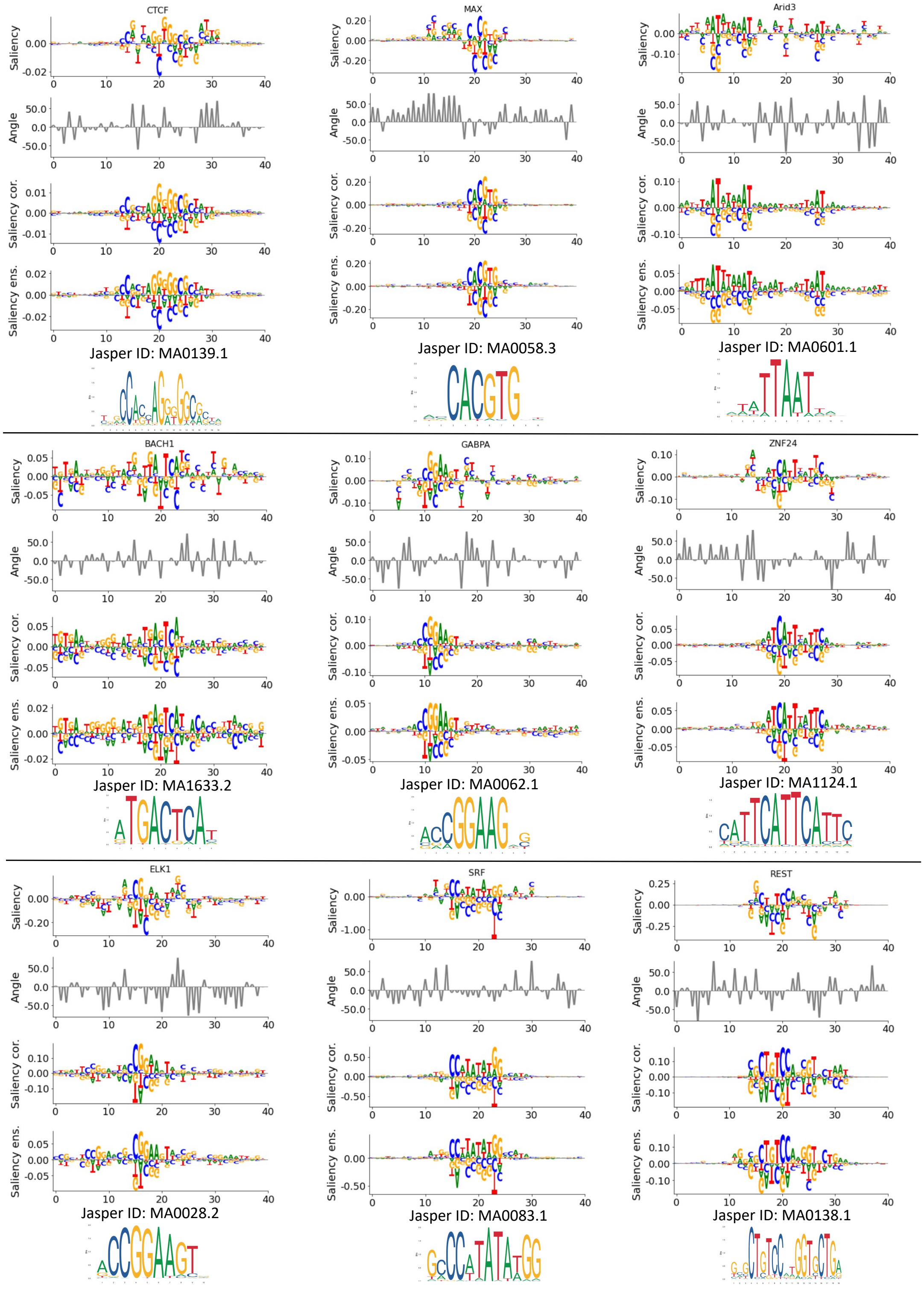


**Supplementary Figure 7.** Saliency map comparison for ChIP-seq data. Comparison of saliency maps before and after correction for CNN-deep-exp for various ChIP-seq experiments: CTCF, MAX, Arid3, BACH1, GABPA, ZNF24, ELK1, SRF and REST. Subplots are representative patches from positive label sequences that show a sequence logo of the saliency scores at each position (top row), a plot of the angles between gradients and the simplex at each position (second row), a sequence logo of the corrected saliency scores (third row), and a sequence logo of the ensemble average (bottom row). Known motifs from JASPAR database are shown below.

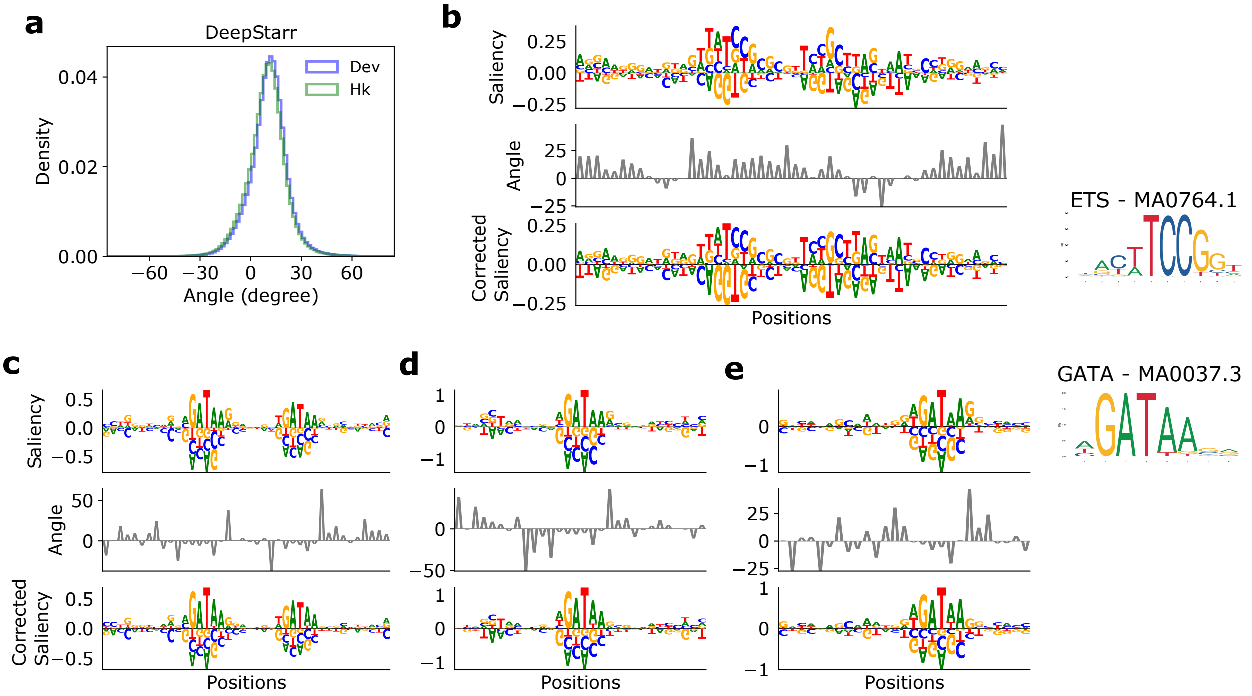


**Supplementary Figure 8.** Gradient correction for DeepSTARR. (a) Histogram of the gradient angles for all test sequences in the DeepSTARR dataset. Different colors represent the different tasks, eg. developmental (Dev) and Housekeeping (HK) enhancer activities. (b-e) Visual comparison of gradient corrections for representative sequences. The sequence logo of the original saliency scores (top row), a plot of the angles between gradients and the simplex at each position (middle row), and a sequence logo of the corrected saliency scores (third row) are shown for a modified DeepSTARR with exponential activations (b) and the original DeepSTARR with ReLU activations (d-e). Known motifs that visually match patterns in the saliency maps are shown on the right labelled with a JASPAR ID.

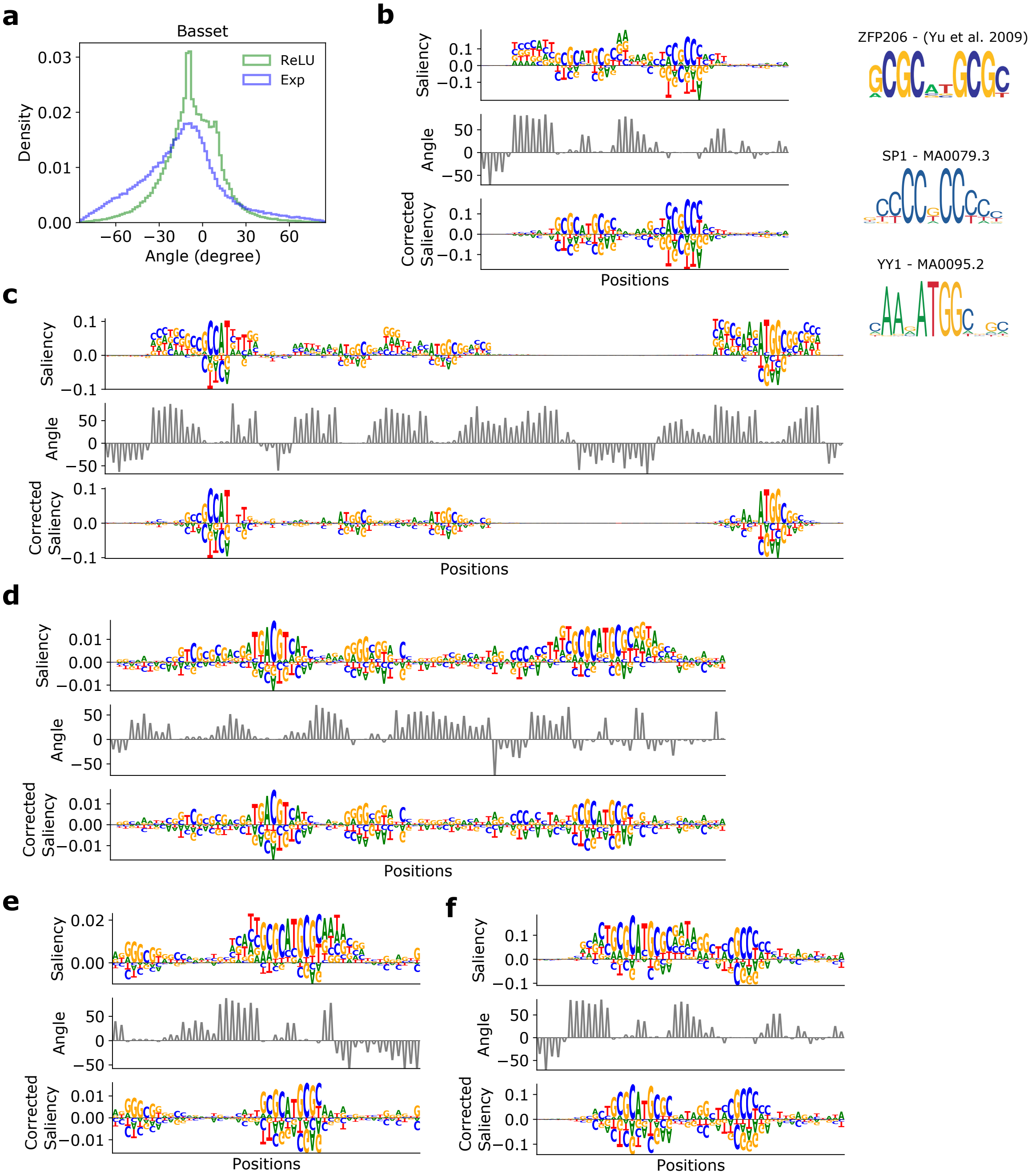


**Supplementary Figure 9.** Gradient correction for Basset. (a) Histogram of the gradient angles for 25,000 test sequences in the Basset dataset, using a Basset model with exponential activations and ReLU activations in first layer filters. (b-f) Visual comparison of gradient corrections for representative sequences. The sequence logo of the original saliency scores at each position (top row), a plot of the angles between gradients and the simplex at each position (middle row), and a sequence logo of the corrected saliency scores (third row) are shown for Basset with exponential activations (b,c) and for ReLU activations (d-f). Known motifs that visually match patterns in the saliency maps are shown on the right labelled with a JASPAR ID.

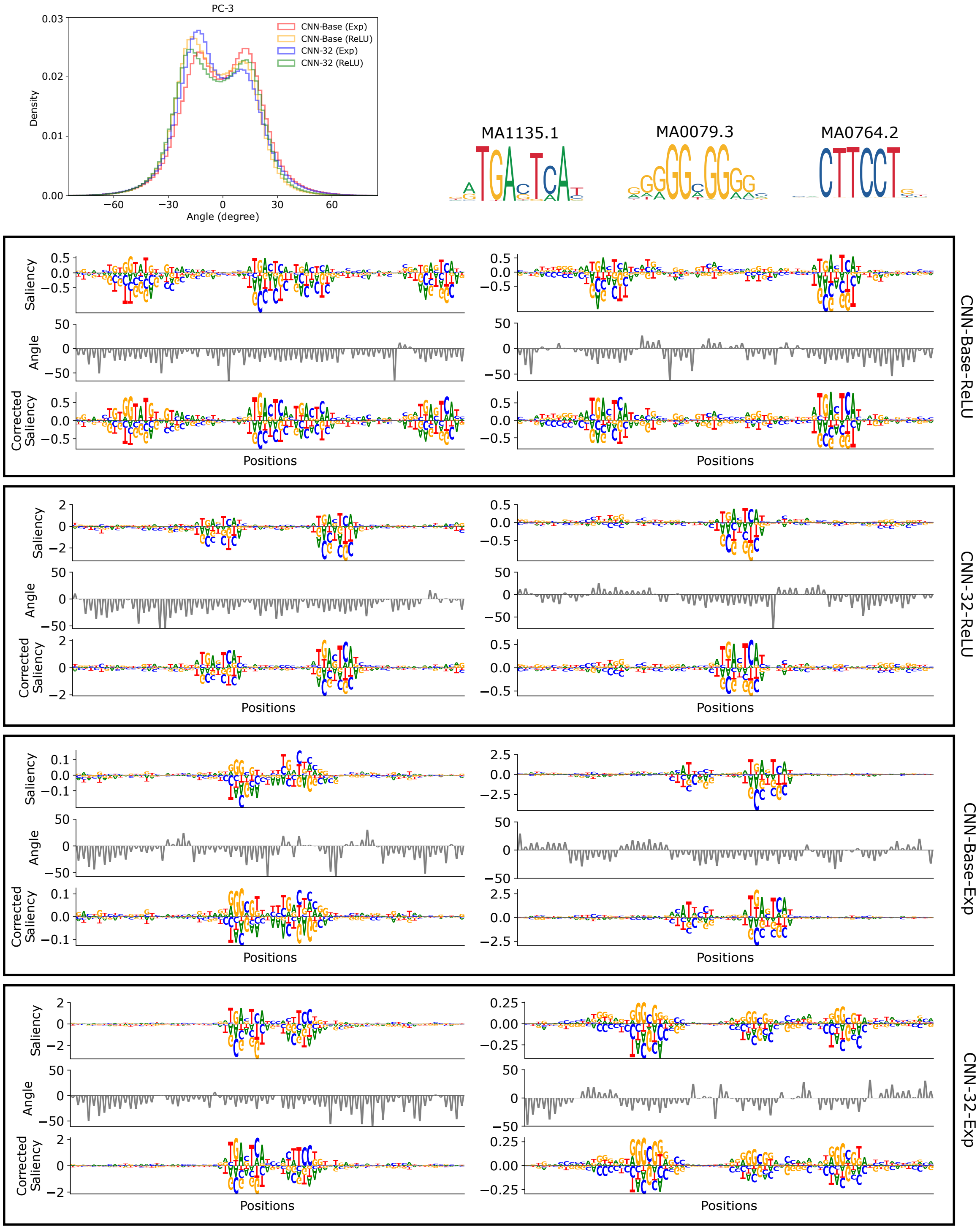


**Supplementary Figure 10.** Gradient correction for quantitative CNNs trained at different resolutions of ATAC-seq data. (Top left) Histogram of the gradient angles of saliency maps from various CNNs (shown in a different color) for all test sequences associated with an IDR peak for PC-3 cell line. (Below) Visual comparison of gradient corrections for two representative sequences for each CNN trained on base- or 32-bin resolution and with ReLU or exponential activations in first layer filters. The sequence logo of the original saliency scores (top row), a plot of the angles between gradients and the simplex at each position (middle row), and a sequence logo of the corrected saliency scores (third row) are shown for two representative sequences for each CNN. Known motifs that visually match patterns in the saliency maps are shown on the top right labelled with a JASPAR ID.

### Supplementary Note 1. Initialization influences extent of off-simplex gradients

We hypothesize that initialization may play a large role in the behavior of the function off of the simplex. To elaborate, if the initialization is set poorly, the initial function may already be pointing far away from the simplex, thereby introducing a larger gradient angle. Since models are trained to minimize the loss, they are only concerned with predictions of observed data (that resides on the simplex). The model then may have limited ability to correct this arbitrary behavior as no data exists off the simplex to fix it during training. To investigate the effect of initialization, we explored how random normal initializations with zero mean and different standard deviations affected the gradient angle of a trained CNN-deep-relu model. In agreement with our hypothesis, we found that the standard deviation in the gradient angle distribution is narrower for smaller initializations and the width of the distribution increases dramatically with larger initializations, with only a marginal drop-off in the classification performance (Supplementary Fig. 11). We noticed a similar trend for other models. This suggests that initialization largely drives the randomness of the function off of the simplex, and it may be beneficial to find a new initialization strategy better suited for categorical inputs such that the initial function better aligns with the simplex.

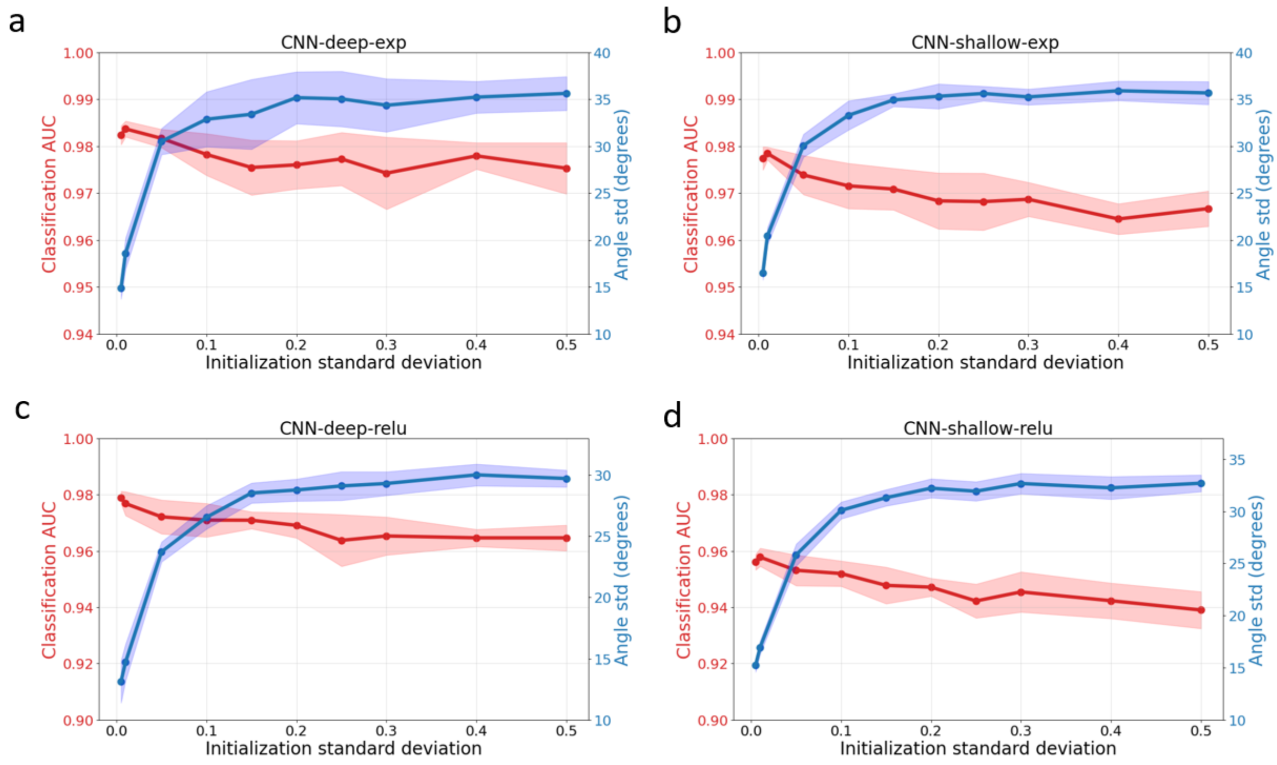


**Supplementary Figure 11.** Initialization analysis using synthetic data. Plot of classification performance (red) and standard deviation of the distribution of gradient angles (blue) for different random normal initializations for CNN-deep-exp, CNN-shallow-exp, CNN-deep-relu and CNN-shallow-relu, trained on synthetic data.

### Supplementary Note 2. Angle regularization during training drives gradients toward simplex

Our proposed correction is ideally suited for post hoc analysis of already trained models. However, it is possible to consider using it as an attribution prior to directly regularize the angle of the input gradients during training to drive the model to actively learn a function that removes this undesirable behavior. To test this, we trained several versions of CNN models with different angle regularization penalties (Supplementary Fig. 12). We found that when sufficient regularization penalty is applied, the gradient angles get driven to zero as expected. We observed a bump in interpretability performance as the angle drops to zero – a small but significant improvement, which is a result of preventing off-simplex gradient noise. Interestingly, the classification performance largely remains constant, until the regularization strength is too large, which then detracts the model from the proper objective. In conclusion, we suggest post-training gradient correction as perhaps a more robust way of correcting the angle noise, as it is simpler to implement (i.e. a single line of code) and does not require carefully tuning an additional hyperparameter.

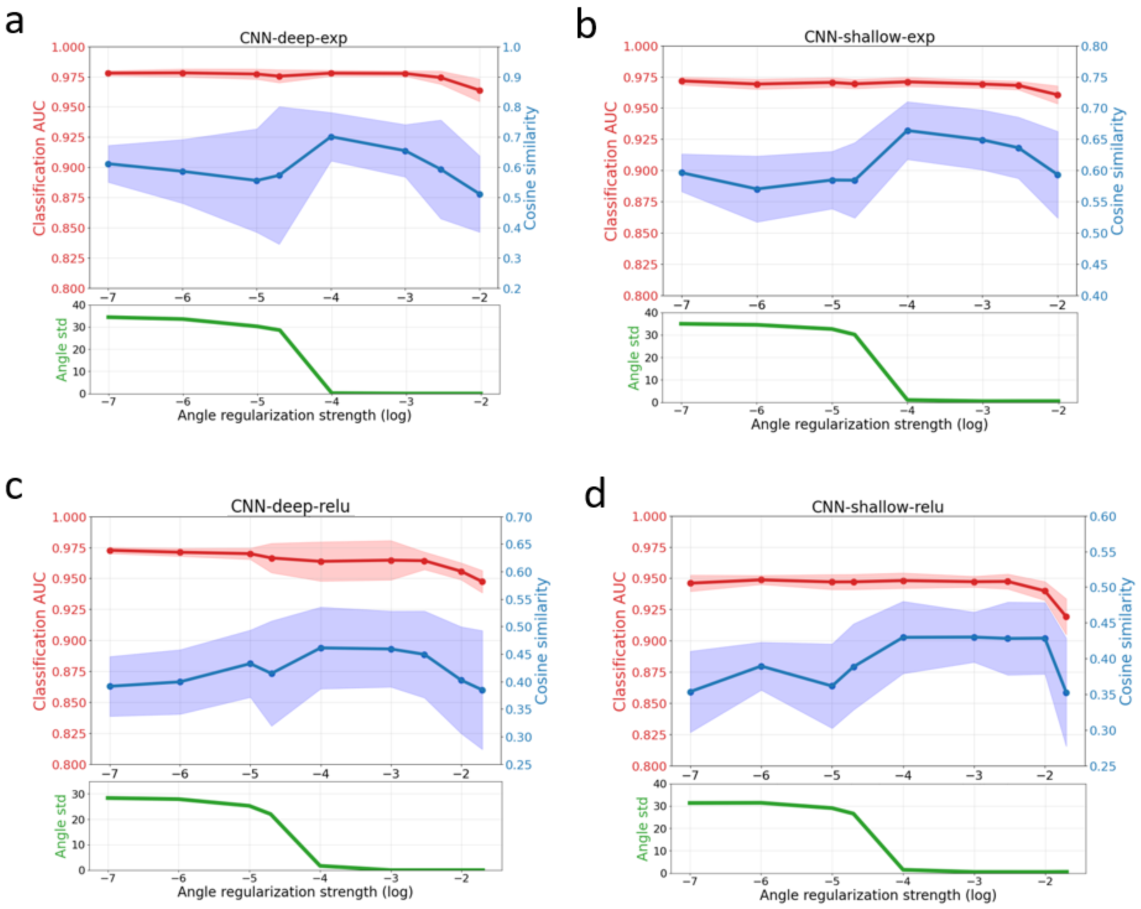


**Supplementary Figure 12.** Regularization analysis using synthetic data. Plot of classification performance (red) and cosine similarity (blue) for different CNNs trained with different gradient angle regularization strengths. Below is a plot of the mean gradient angle distribution for models trained with each regularization strength. Dots represent the average across 10 trials and the shaded region represents the standard deviation of the mean. Additional details: We regularized the angle of the input gradients during training by penalizing the mean input angle for each batch with a hyperparameter (which is varied along the $x$-axis). We trained 5 CNN models for each regularization penalty setting using different random initializations. When sufficient regularization penalty is applied, the gradient angles get driven to zero as expected.
